# An evolutionarily conserved leucine zipper-like motif accounts for strong tetramerization capabilities of SEPALLATA-like MADS-domain transcription factors controlling flower development

**DOI:** 10.1101/125294

**Authors:** Florian Rümpler, Günter Theißen, Rainer Melzer

## Abstract

The development of angiosperm flowers is regulated by homeotic MIKC-type MADS-domain transcription factors that activate or repress target genes via the formation of DNA-bound, organ specific tetrameric complexes. The protein-protein interaction (PPI) capabilities differ considerably between different MIKC-type proteins. The floral homeotic protein SEPALLATA3 (SEP3) acts as a hub that incorporates numerous other MADS-domain proteins into tetrameric complexes that would otherwise not form. However, the molecular mechanisms that underlie these promiscuous interactions remain largely unknown. In this study we created a collection of amino acid substitution mutants of SEP3 to quantify the contribution of individual residues on protein tetramerization during DNA-binding, employing methods of molecular biophysics. We show that leucine residues at certain key positions form a leucine zipper structure that is essential for tetramerization of SEP3, whereas the introduction of physicochemically very similar residues at respective sites impedes the formation of DNA-bound tetramers. Comprehensive molecular evolutionary analyses of MADS-domain proteins from a diverse set of flowering plants revealed exceedingly high conservation of the identified leucine residues within SEP3-subfamily proteins throughout angiosperm evolution. In contrast, MADS-domain proteins that are unable to tetramerize among themselves exhibit preferences for other amino acids at homologous sites. Our findings indicate that the subfamily-specific conservation of amino acid residues at just a few key positions account for subfamily-specific interaction capabilities of MADS-domain transcription factors and shaped the present-day structure of the PPI network controlling flower development.

## INTRODUCTION

Complexity of biological systems is often achieved by the combinatorial activity of a small number of factors (Remenyi, et al. 2004). One important example are protein-protein interaction (PPI) networks that are based on transcription factors (TFs) that act in a combinatorial manner to accomplish the required degree of e.g. morphological complexity. PPI networks often approximate a scale-free structure (Barabasi and Oltvai 2004). They contain a small number of hub proteins with many interaction partners and a large number of poorly connected nodes. Though combinatorial control is of eminent importance for almost all developmental processes, the molecular determinants that are underlying the specific combinatorial interactions remain poorly understood. This is especially true for protein-protein interactions among TFs belonging to the same family. The respective TFs are often very similar in terms of sequence and biochemical properties yet fulfill highly distinct and specific functions which are at least partially determined by distinct protein-protein interactions. The PPI network controlling flower development in angiosperms is a good case in point. Floral organ specification is regulated by so called floral quartets which are organ specific tetrameric complexes of MIKC-type MADS-domain TFs bound to two adjacent DNA-binding sites while looping the DNA to regulate target genes (Coen and Meyerowitz 1991; Pelaz, et al. 2000; Theißen and Saedler 2001; Ditta, et al. 2004; Melzer and Theißen 2009; Melzer, et al. 2009). In the model plant species *Arabidopsis thaliana* the floral homeotic protein SEPALLATA3 (SEP3) together with its paralogs SEP1, SEP2 and SEP4 from the closely related LOFSEP-subfamily bears a central role by forming tetrameric complexes with numerous other MIKC-type MADS-domain TFs (Zahn, et al. 2005; Immink, et al. 2009; Melzer and Theißen 2009; Smaczniak, et al. 2012). The four SEP proteins act in a largely redundant manner but in agreement with their central position in the PPI network controlling flower development *sep* multiple mutants show severe developmental defects (Pelaz, et al. 2000; Ditta, et al. 2004). *sep1 sep2 sep3* triple mutant plants develop sepals from primordia that would normally develop into petals, stamens and carpels and *sep1 sep2 sep3 sep4* quadruple mutants develop vegetative leaves instead of floral organs (Pelaz, et al. 2000; Ditta, et al. 2004).

Among the four SEP genes, SEP3 has been studied best (Favaro, et al. 2003; Immink, et al. 2009; Melzer and Theißen 2009; Melzer, et al. 2009; Smaczniak, et al. 2012; Jetha, et al. 2014). Beyond the formation of complexes that determine floral organ identity SEP3 is also involved in controlling flowering time, floral transition and ovule development (Favaro, et al. 2003; Immink, et al. 2009; Liu, et al. 2009; Lopez-Vernaza, et al. 2012). It does therefore constitute one of the major hub proteins within the PPI network controlling reproductive development (Favaro, et al. 2003; Immink, et al. 2009; Liu, et al. 2009; Smaczniak, et al. 2012). However, it is unclear which biochemical and biophysical properties enable SEP3 to form DNA bound tetramers with numerous partners whereas other MIKC-type MADS-domain TFs are unable to form floral quartet like complexes among themselves. For example, the floral homeotic proteins APETALA3 (AP3) and PISTILLATA (PI) from *A. thaliana* that are involved in the developmental specification of petals and stamens do only form obligate heterodimers and require SEP proteins for tetramer formation (Immink, et al. 2009; Melzer and Theißen 2009; Melzer, et al. 2014).

The protein-protein interactions that allow for tetramer formation are mainly mediated by the about 80 amino acids long keratin-like domain (K-domain), which is shared by all MIKC-type MADS-domain TFs (Yang, et al. 2003; Yang and Jack 2004; Melzer and Theißen 2009). The amino acid sequence within the K-domain of most MADS-domain proteins shows three characteristic heptad repeat patterns (K1-; K2-; K3-subdomain repeat) of the form [abcdefg]_n_, where most ‘a’ and ‘d’ positions are occupied by highly hydrophobic residues (Riechmann and Meyerowitz 1997; Yang, et al. 2003; Yang and Jack 2004). This sequence feature is typical for coiled-coils, a common and intensively studied type of protein-protein interaction domains (Betz, et al. 1995; Mason and Arndt 2004; Parry, et al. 2008; Mason, et al. 2009) (Figure 1). Within a coiled-coil, an α-helix is formed and the amino acids on heptad repeat ‘a’ and ‘d’ positions form a stripe of hydrophobic residues that runs along the α-helix and facilitates hydrophobic interaction with a partner coiled-coil (Mason and Arndt 2004; Mason, et al. 2009).

**Figure 1.**
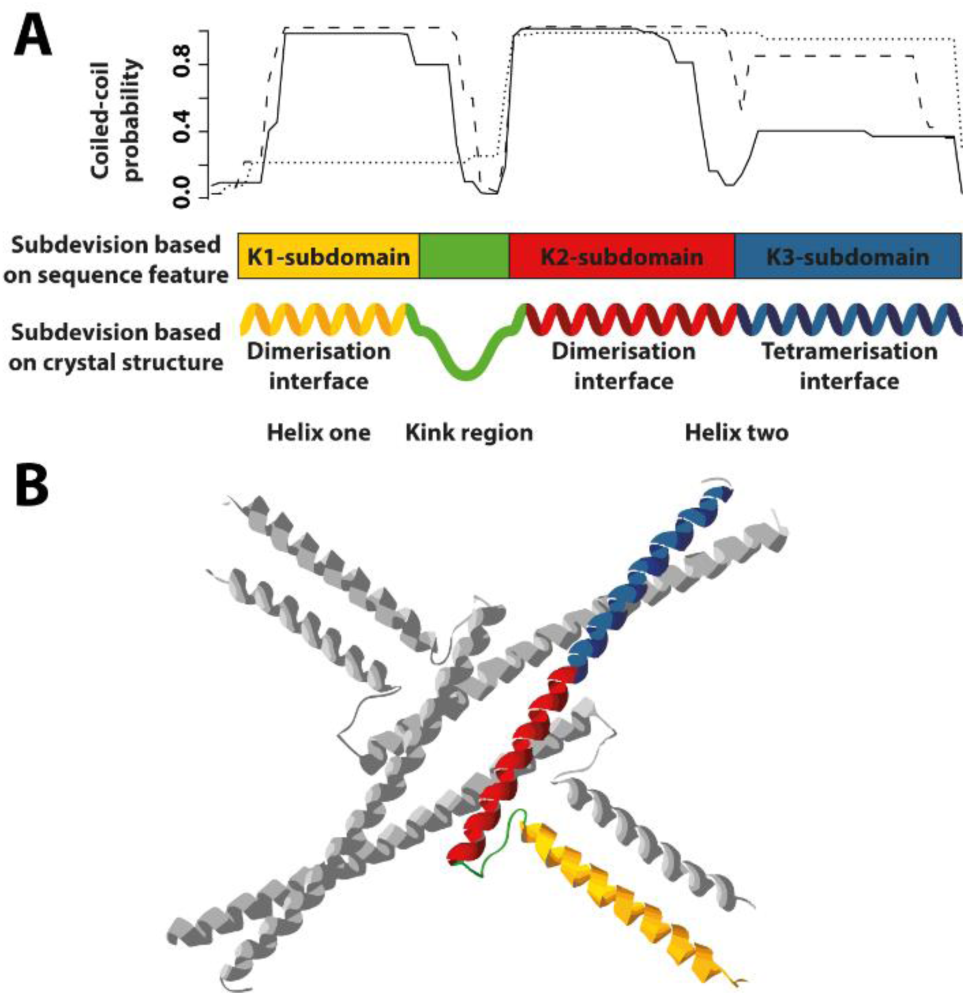
Domain architecture of the K-domain of SEP3 based on sequence and structural features. (**A**) Based on coiled-coil predictions (top) the K-domain was assumed to fold into three separate coiled-coils and was thus subdivided into three subdomains K1-, K2- and K3-subdomain (middle). The crystal structure of the K-domain of SEP3 revealed that it folds into two α-helices separated by a kink region (bottom). The first helix spans the K1-subdomain (color coded in yellow) and is involved in the dimerisation of two SEP3 monomers (i.e. dimerisation interface). The second helix spans the K2- and K3-subdomains and constitutes an N-terminal interaction interface that further stabilizes dimerisation of two SEP3 monomers (red) and a second C-terminal interaction interface that mediates the interaction of two SEP3 dimers (i.e. tetramerisation interface, blue). Coiled-coil predictions were performed with COILS (Lupas, et al. 1991). The solid, dashed and dotted lines correspond to a sliding window size of 14, 21 and 28 amino acids used for the prediction, respectively. (**B**) Crystal structure of a SEP3 K-domain homotetramer (Puranik, et al. 2014). The dimerisation interface of helix one, the kink region, the dimerisation interface of helix two and the tetramerisation interface of one K-domain are color coded in yellow, green, red and blue, respectively.

Recently the crystal structure of the complete K-domain of SEP3 was reported (Puranik, et al. 2014). Based on the crystal structure, the K-domain forms two amphipathic α-helices separated by a kink region which prevents intramolecular association of both helices. Helix one comprises the first heptad repeat (K1-subdomain) and is involved in dimerisation of two SEP3 monomers. Helix two spans heptad repeat two (K2-subdomain) that further stabilizes the interaction of two SEP3 monomers and heptad repeat three (K3-subdomain) which constitutes an interface for the interaction of two SEP3 dimers i.e. tetramerisation (Figure 1). In this study, we determine sequence features that enable SEP3 to form tetrameric complexes and identify the amino acid patterns that distinguish members of the SEP3 subfamily from other MIKC-type MADS-domain TFs with more restricted tetramerization capabilities. Our data suggest that leucine residues at intramolecular contact points and at the interaction interface of the K3-subdomain are indispensable for tetrameric complex formation. Sequence analyses of MIKC-type MADS-domain TFs from a broad set of flowering plant species reveal very high conservation of the examined leucine residues in the SEP3 subfamily throughout angiosperm evolution. In contrast, members of other MIKC-type MADS-domain TF subfamilies such as AP3 and PI show preferences for other amino acids at homologous sites. The identified leucines may thus be a critical denominator that determines the ability of SEP3-subfamily proteins to incorporate a great number of other MIKC-type MADS-TFs into floral quartets.

## RESULTS

### Leucine residues in the K-domain strongly influence cooperative DNA-binding of SEP3

To investigate the relevance of the different K-subdomains for cooperative DNA-binding and tetramer formation, single and double amino acid substitutions to proline were performed. Proline was chosen because it possesses helix-breaking properties (Richardson 1981; Nilsson, et al. 1998) and would thus be expected to disrupt the overall structure of the respective K-subdomain. For each of the three K-subdomains two substitution mutants were created (Figure 2A, Supplementary Figure S1). Based on coiled-coil predictions one substitution mutant was supposed to destroy the K-subdomain coiled-coil (L115P for K1-, L131P-L135P for K2- and L164P for K3-subdomain, respectively) whereas the other one was expected to not alter the formation of the respective coiled-coil (S94P (K1); L145P (K2); G178P (K3)). Beyond the three K-subdomains, we also introduced proline substitutions at positions occupied by two conserved hydrophobic amino acids in the interhelical region between the K1- and the K2-subdomain (L120P and L123P, Figure 2A) because positions homologous to L120 and L123 have been shown to be important for the interaction of MADS-domain proteins (Yang and Jack 2004).

**Figure 2.**
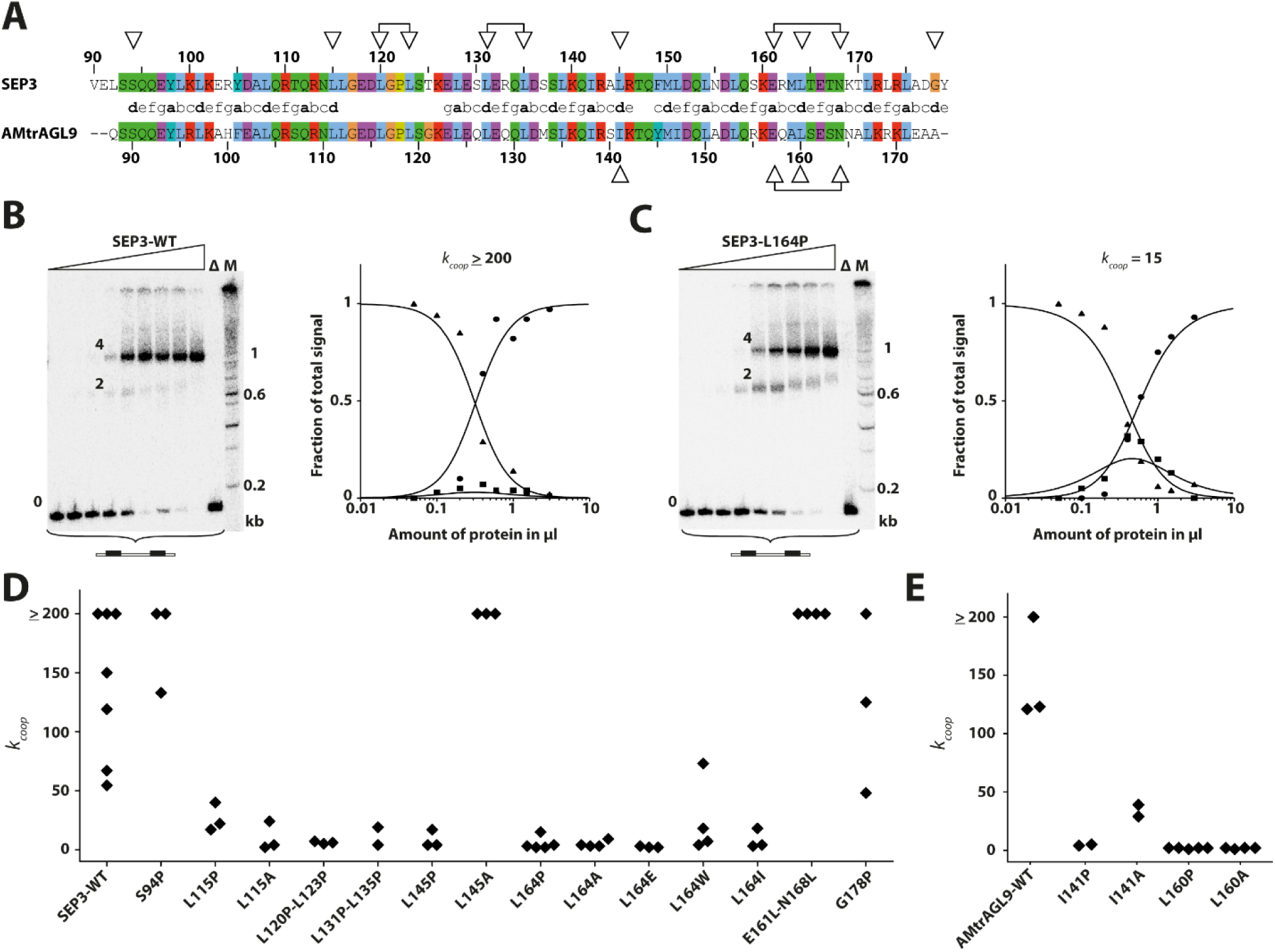
Ability of SEP3 and AMtrAGL9 wild type proteins and different amino acid substitution mutants to cooperatively bind to DNA. (**A**) Pairwise sequence alignment of the K-domains of SEP3 and AMtrAGL9. The heptad repeat pattern is depicted in the center. Positions at which amino acids were substituted are indicated by open triangles. (**B** and **C**) Binding of SEP3 wild type (**B**) and SEP3-L164P (**C**) to a DNA probe containing two CArG-boxes. Increasing amounts of *in vitro* translated protein were incubated with constant amounts of DNA probe. As negative control the empty pTNT vector without any cDNA insert was used as template DNA for the *in vitro* translation (lane ∆). For size comparison a radioactively labeled DNA ladder (100 bp Ladder, NEB) was applied (lane M). The labeling of the three different fractions ‘0’, ‘2’ and ‘4’ corresponds to the number of proteins bound to one DNA molecule. Quantified signal intensities of the different fractions and graphs, fitted according to equation (1) to (3) described in Methods, are shown next to the gel pictures (triangle: free DNA; square: DNA probe bound by two proteins; circle: DNA probe bound by four proteins). The *k*_*coop*_ value inferred from this particular measurement is depicted above the diagram. (**D** and **E**) *k*_*coop*_ values for the wild type protein and all examined single and double amino acid substitution mutants of SEP3 (**D**) and AMtrAGL9 (**E**). *k*_*coop*_ values above 200 could not be determined reliably (see Methods).

We used electrophoretic mobility shift assays (EMSAs) to study the DNA-binding and tetramerisation behavior of the mutant SEP3 proteins. Based on previous studies it is known that SEP3 binds as homodimer to a DNA-element termed CArG-box (for CCArichGG; consensus sequence 5’-CC(A/T)6GG-3’) and that four SEP3 proteins bind to a DNA probe containing two CArG-boxes (Melzer, et al. 2009). To first investigate whether DNA-binding affinities of individual dimers were affected by the different amino acid substitutions, we performed saturation binding EMSA experiments using increasing amounts of a DNA probe containing only one CArG-box together with constant amounts of protein as previously described (Jetha, et al. 2014). The estimated affinities for binding of the altered SEP3 proteins to a single DNA-binding site varied slightly but did not considerably differ from the values obtained for SEP3 wild type protein (Supplementary Figure S2, Supplementary Table S1), indicating that the different amino acid substitutions did not or only marginally affect DNA-binding of individual dimers.

If increasing amounts of SEP3 were incubated together with constant amounts of a DNA probe containing two CArG-boxes, three bands of different electrophoretic mobility were observed (Figure 2B left side). As determined previously (Melzer, et al. 2009) the band of high electrophoretic mobility constitutes unbound DNA (indicated with ‘0’ in Figure 2B), the band of intermediate electrophoretic mobility constitutes a DNA probe bound by two SEP3 proteins (‘2’) and the band of low electrophoretic mobility constitutes a DNA probe bound by four SEP3 proteins (‘4’). By analyzing the signal intensities of the three different fractions the ability of SEP3 to form DNA-bound tetrameric complexes can be quantified and expressed via the cooperativity constant *k*_*coop*_ (equation (4) in Methods). *k*_*coop*_ equals 1 for non-cooperative binding and increases with increasing tetramer formation capabilities of the examined protein. SEP3 wild type protein always showed a highly cooperative DNA-binding although the degree of cooperativity varied between different experiments and was slightly higher than previously estimated (Melzer, et al. 2009; Jetha, et al. 2014), probably owing to difficulties to precisely determine high *k*_*coop*_ values (Figure 2B and D, Supplementary Table S1).

In contrast to the wild type protein, all of the leucine-to-proline substitution mutants of SEP3 (L115P; L120P-L123P; L131P-L135P; L145P; L164P) showed a considerably reduced ability to bind cooperatively to DNA *in vitro*, independent of whether the formation of coiled-coils was predicted to be affected or not (Figure 2C and D, Supplementary Table S1). Only the two proline substitutions S94P and G178P, located at the N- and C-terminal borders of the K-domain, respectively, did not strongly reduce cooperative binding of SEP3.

To test the effect of amino acid substitutions that are supposed to have a less severe effect on helix formation than proline, we substituted a subset of the previously selected leucines (L115; L145; L164) by alanine. Surprisingly, of these 3 substitutions only L145A showed a cooperative DNA-binding ability comparable to that of SEP3 wild type protein, whereas substitutions L115A and L164A caused an almost complete loss of cooperative DNA-binding, comparable to the proline substitutions at the respective positions (Figure 2D, Supplementary Table S1). We further substituted position L164 by three additional amino acids (L164E; L164W; L164I) comprising glutamate and tryptophan which occur at position 164 in several members of the SEP subfamily and isoleucine which has very similar physicochemical properties to leucine. However, none of the resulting mutants was able to approach SEP3 wild type cooperative binding strength (Figure 2D, Supplementary Table S1). Our results indicate that the examined leucine residues are of critical importance for tetramer formation and cooperative binding of SEP3.

Within the [abcdefg]_n_ heptad repeat of the K3-subdomain of SEP3 two neighboring ‘a’ positions (E161; N168) are not occupied by hydrophobic amino acids. Substituting these positions by leucine (E161L-N168L) resulted in a higher probability for the formation of the K3-subdomain coiled-coil *in silico* (Supplementary Figure S1). The respective mutant protein showed a cooperativity at least as high as the wild type protein in EMSAs. In contrast to the wild type protein, repeated measurements yielded *k*_*coop*_ values that consistently were above 200 (Figure 2D, Supplementary Table S1). In fact, in none of the performed EMSAs a signal of a DNA probe bound by only one protein dimer was detected, an observation that was different from the other proteins for which high cooperativity in DNA-binding was detected (e.g. SEP3-WT and SEP3-L145A) indicating that cooperative binding was increased by the E161L-N168L substitutions (Supplementary Figure S3). Surprisingly, when we performed saturation binding EMSA experiments using increasing amounts of a DNA probe containing only one CArG-box the mutant protein SEP3-E161L-N168L exhibited no binding of individual dimers. Instead a signal of low electrophoretic mobility occasionally occurred for high amounts of applied DNA probe that might constitutes a protein DNA complex consisting of more than two proteins (Supplementary Figure S4).

### Mutations in the most distantly related ortholog of SEP3 have very similar effects on cooperative DNA-binding as in SEP3

Next we aimed to assess whether the importance of the identified leucine residues is evolutionarily conserved within the SEP3-subfamily. The MADS-domain TF AMtrAGL9 from the earliest diverging angiosperm *Amborella trichopoda* constitutes the most distantly related ortholog of SEP3 (Zahn, et al. 2005). In EMSA experiments AMtrAGL9 forms homotetrameric protein-DNA complexes with a cooperative binding affinity comparable to SEP3 (Figure 2E, Supplementary Figure S5). AMtrAGL9 amino acid position I141 is homologous to SEP3 L145 and is thus located in the K2-subdomain heptad repeat of AMtrAGL9 (Figure 2A). Substitution to alanine at that position interfered to some extent with cooperative binding capabilities whereas substitution to proline at position I141 results in an almost complete loss of cooperative binding (Figure 2E, Supplementary Table S1). If amino acid position L160 of AMtrAGL9, which is homologous to position L164 in the center of the K3-subdomain of SEP3, is exchanged by proline or alanine, the ability of AMtrAGL9 to cooperatively bind to DNA is almost completely lost in either case, a behavior that is similar to that observed for SEP3 (compare Figure 2D and E).

### Interacting sites are more often occupied by leucine in SEP3-subfamily proteins than in proteins of other MIKC-type subfamilies

The importance of leucine residues for the tetramerisation ability of SEP3 and AMtrAGL9 raised the question as to which extent these positions are conserved within the SEP3 subfamily and which amino acid preferences members of other MIKC-type protein subfamilies show at homologous sites. We therefore created a multiple sequence alignment based on 1,325 sequences of MIKC-type MADS-domain proteins belonging to 14 subfamilies and comprising sequences from a diverse array of seed plants. Despite the high evolutionary distance of the sampled taxa, the sequences aligned almost without gaps throughout the complete K-domain (i.e. without potential insertions or deletions). The only exception were PI-subfamily protein sequences, among which a deletion of four amino acids within the C-terminal half of the K-domain was very common. This deletion within the PI-linage most likely occurred after early diverging angiosperms branched off, as most of the sampled PI-subfamily sequences from early diverging angiosperms still possess those four amino acids.

We first compared the conservation of sites that are homologous to the 15 residues that (based on the crystal structure of SEP3) mediate the hydrophobic intra- and intermolecular interactions in the SEP3 homotetramer (Puranik, et al. 2014) to the overall conservation of the K-domain. We found that within the SEP3 subfamily, sites that are homologous to interacting sites in the SEP3 homotetramer are significantly less variable than the remaining residues of the K-domain (Figure 3A). This conservation pattern also holds true for sequences of all other 13 subfamilies of MIKC-type MADS-domain proteins (Figure 3B, Supplementary Figure S6A) as well as for sequences from gymnosperms to core eudicots (Supplementary Figure S6B). Beyond this similar pattern of conserved positions also the amino acid properties in terms of hydrophobicity at homologous sites appear highly similar among all examined subfamilies (Supplementary Figure S7), suggesting that the overall structure of the K-domain as determined for SEP3 is conserved among MIKC-type proteins of most if not all subfamilies and throughout seed plants.

**Figure 3.**
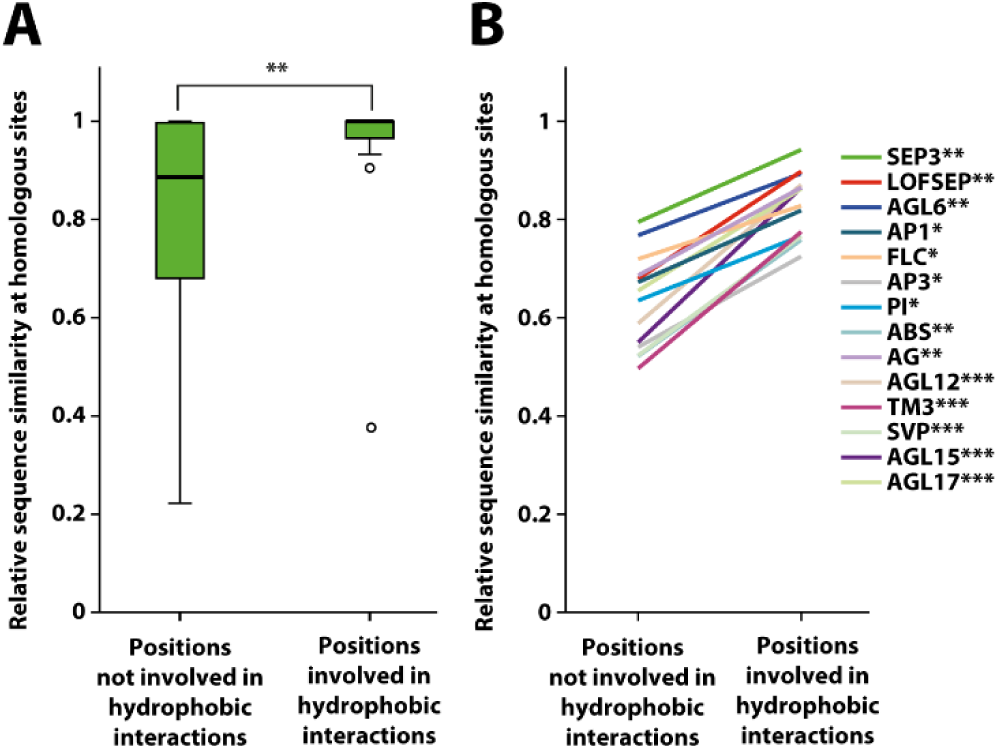
Sequence similarity analysis of SEP3-subfamily proteins and members of other MIKC-type MADS-domain protein subfamilies. (**A**) Box-plot showing relative sequence similarity at homologous sites of SEP3-subfamily proteins for positions that are involved in hydrophobic interactions within the SEP3 homotetramer and positions that are not involved in hydrophobic interactions. (**B**) Line graph showing the same analysis as in (**A**) but for all MIKC-type protein subfamilies. For all subfamilies amino acid positions that are homologous to sites involved in hydrophobic interactions are significantly less variable than positions that are homologous to non-interacting sites (Mann-Whitney-U-test; * p = 0.01-0.05; ** p = 0.001-0.01; *** p < 0.001).

Next we analyzed the amino acid distribution at sites homologous to the 12 leucine residues (L101, L108, L115, L120, L123, L128, L131, L135, L154, L157, L164 and L171) that contribute to inter- and intramolecular interactions in a SEP3 homotetramer (Figure 4A) (Puranik, et al. 2014). Notwithstanding the high evolutionary distance of the examined SEP3-subfamily proteins all these leucine residues were found to be highly conserved within the 78 examined sequences; 8 out of 12 positions were completely invariable (Figure 4B). In contrast to this, members of other subfamilies (e.g. AP3- and PI-subfamily proteins) often show preferences for other amino acids on equivalent sites (Figure 4B, Supplementary Figure S8). Especially positions equivalent to L154, L157 and L164 of SEP3 that are located within the center of the tetramerisation interface are often not occupied by leucines in AP3- and PI-subfamily proteins. The high even though slightly lower conservation of leucines also becomes apparent within LOFSEP-subfamily proteins (comprising SEP1, SEP2 and SEP4 from *A. thaliana*) which form the sister group of SEP3-subfamily proteins and that are assumed to function in a mostly redundant manner with SEP3 during flower development (Figure 4C) (Pelaz, et al. 2000; Ditta, et al. 2004; Zahn, et al. 2005). The closest relatives of SEP3- and LOFSEP-subfamily proteins are AGL6-subfamily proteins followed by AP1-subfamily proteins (Kim, et al. 2013). However, despite the close relationship AGL6- as well as AP1-subfamily proteins display a considerably lower leucine frequency especially on sites within the tetramerisation interface (Figure 4C, Supplementary Figure S8). Instead, these positions are more frequently occupied by other hydrophobic amino acids such as isoleucine and methionine. It has previously been shown that within a coiled-coil, leucine packs very well at heptad repeat ‘d’ positions and enables the formation of a tight dimeric coiled-coil as it becomes apparent in a leucine-zipper (Zhu, et al. 1993; Betz, et al. 1995). In contrast other hydrophobic amino acids such as isoleucine or valine lead to steric hindrance at heptad repeat ‘d’ positions (Betz, et al. 1995) (Figure 4D).

**Figure 4.**
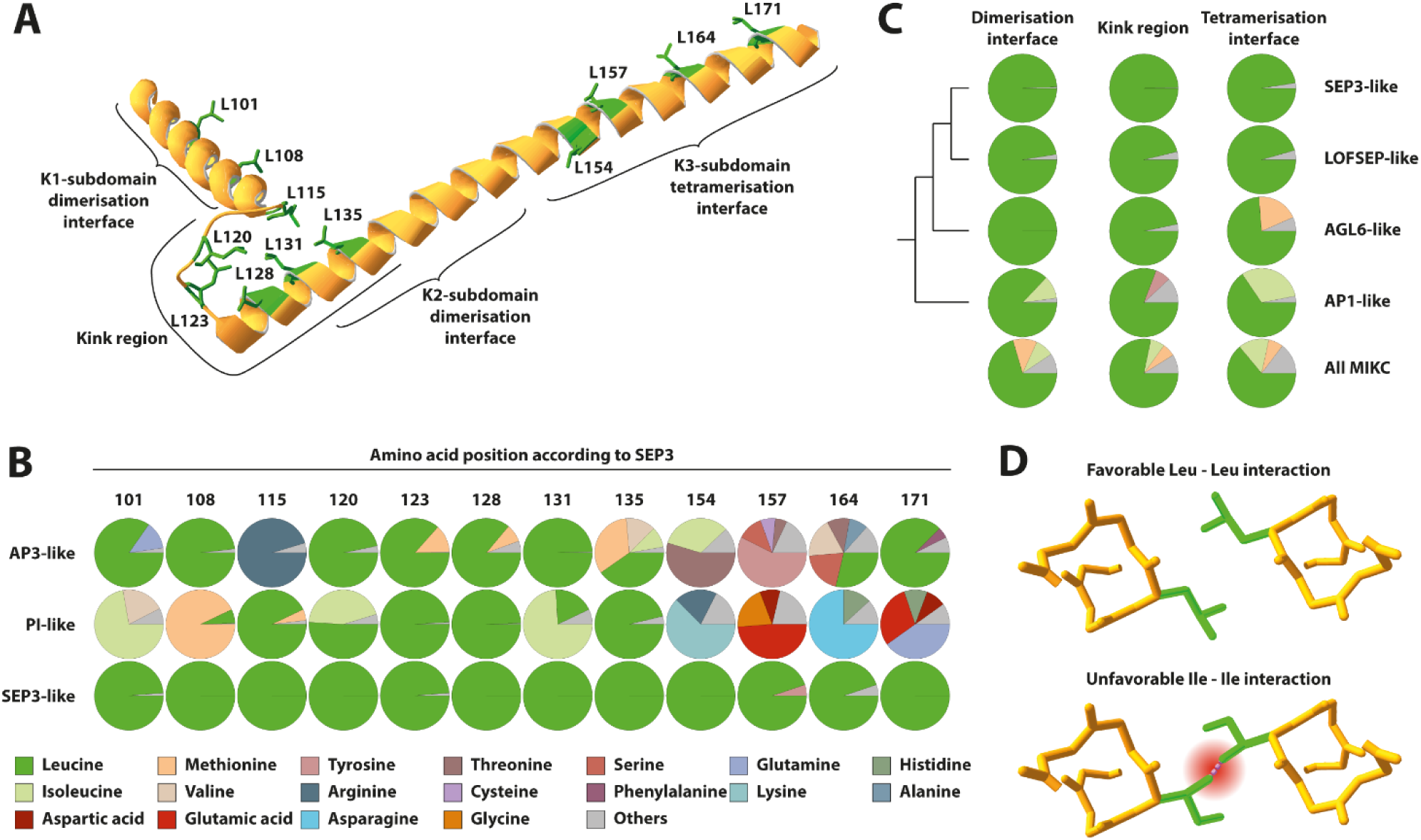
Amino acid preferences of SEP3-subfamily proteins and members of other MIKC-type MADS-domain protein subfamilies. (**A**) Picture of the crystal structure of a single K-domain of SEP3. Leucine side chains that are involved in inter- and intramolecular interactions are shown in green. (**B**) Amino acid frequencies at sites homologous to leucine residues that are involved in inter- and intramolecular interactions in the SEP3 homotetramer shown for SEP3-, AP3- and PI-subfamily proteins. Amino acids that occurred in less than 5 % of the examined subset of sequences were condensed as ‘others’. The vast majority of the positions shown vertically are homologous to each other. The only exception are positions 154, 157, 164 and 171 of PI-like proteins. In this case, a gap was detected in the alignment but amino acids directly following the gap were included here. (C) Amino acid preferences at sites homologous to leucine residues that contribute to dimerisation interface (L101, L108), kink region (L115, L120, L123, L128, L131, L135) and tetramerisation interface (L154, L157, L164, L171) in the SEP3 homotetramer, shown for SEP3-, LOFSEP-, AGL6- and AP1-subfamily proteins and all MIKC-type proteins, respectively, following the color coding of panel B. (**D**) Part of the crystal structure of two interacting tetramerisation interfaces (D) within a SEP3 homotetramer. The picture illustrates the favorable Leu-Leu interaction at heptad repeat ‘d’ positions as it becomes apparent several times within a SEP3 homotetramer (upper part). In contrast to the γ-branched leucine a β-branched amino acid such as isoleucine would potentially lead to steric hindrance at heptad repeat ‘d’ positions (lower part).

### Insertion of leucine residues into the K3-subdomain of AP3 facilitates homotetramerisation of the chimeric protein SEP3AP3chim

Based on our data we hypothesized that the overall structure of the K-domain is conserved throughout most if not all subfamilies of MIKC-type MADS-domain proteins and that evolutionarily conserved preferences for different amino acids on interacting sites account for subfamily-specific interaction capabilities. We aimed to test our hypothesis with help of the chimeric protein SEP3_AP3chim_, in which we substituted the K3-subdomain (i.e. tetramersiation interface) of SEP3 (residues 150-181) by the homologous sites of AP3 (Figure 5A and B). AP3 is known to form obligate heterodimers with PI and is thus not able to form DNA-binding homodimers or homotetramers (Moitra, et al. 1997; de Folter, et al. 2005). As expected, the chimeric protein SEP3_AP3chim_ showed a complete loss of homotetramerisation capabilities compared to SEP3 wild type protein in EMSA experiments (Figure 5C and D right side, Supplementary Table S1). Although the K3-subdomains of SEP3 and AP3 share only four identical residues at homologous sites the sequence similarity in terms of hydrophobicity on most heptad repeat ‘a’ and ‘d’ positions is comparatively high (Figure 5A). However, two heptad repeat ‘d’ positions occupied by leucine in SEP3 (L157 and L164) are occupied by threonine and glutamine in AP3, respectively (Figure 5A). Both leucines are highly conserved throughout SEP3-subfamily proteins whereas homologous sites in AP3-subfamily proteins are almost exclusively occupied by residues other than leucine (Figure 4B). We thus substituted positions T157 and Q164 of the chimeric protein by leucine and tested the ability of the resulting mutants to form homotetramers. Both single amino acid substitutions could not improve tetramerisation ability of the chimeric protein (Supplementary Table S1). However, the insertion of both leucine residues into the K3-subdomain of SEP3_AP3chim_ sufficed to fully ‘restore’ the ability to form DNA-binding homotetramers (Figure 5E right side). Visualizing the amino acid sequence of the tetramerisation interface of SEP3 and AP3 in a helical wheel diagram illustrates how residues M150, L157, L164 and L171 form a strong hydrophobic stripe within the tetramerisation interface of SEP3, whereas the hydrophobic stripe is interrupted by threonine and glutamine in AP3 (Figure 5C and D left side). Substituting both residues by leucine closes the gap within the hydrophobic stripe and most likely thereby facilitates homotetramerisation (Figure 5E left side).

**Figure 5.**
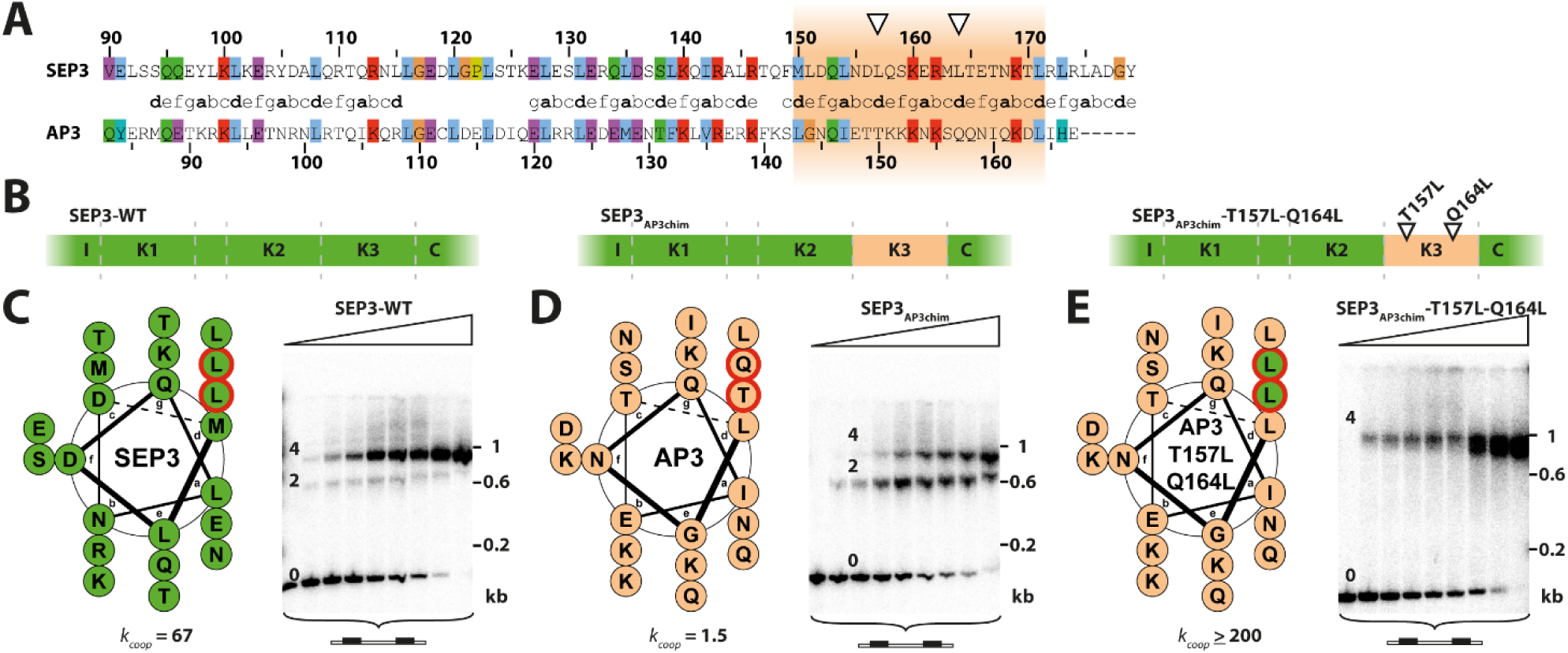
Design and cooperative DNA-binding capabilities of the chimeric protein SEP3_AP3chim_. (**A**) Pairwise sequence alignment of the K-domains of SEP3 and AP3. The heptad repeat pattern is depicted in the center. Orange background marks the region that was substituted to create the chimeric protein SEP3_AP3chim_. Open triangles mark the positions of the subsequently introduced amino acid substitutions. (**B**) Experimental setup to test for the ability of leucines to restore tetramerisation ability of SEP3_AP3chim_. First the complete tetramerisation interface (i.e. the K3-subdomain) of SEP3 was substituted by the equivalent positions of AP3. Subsequently the two residues T157 and Q164 were substituted back to leucine to reestablish the hydrophobic stripe. (**C**-**E**, left) Helical wheel diagram of the tetramerisation interface of SEP3 wild type (**C**), SEP3_AP3chim_ (E) and SEP3_AP3chim_-T157L-Q164L (**E**), respectively, to illustrate the presumed position of amino acids 157 and 164 (framed in red) within the hydrophobic stripe of the K3-subdomain coiled-coil. (**C**-**E**, right) Binding of SEP3 wild type (**C**), SEP3_AP3chim_ (**D**) and SEP3_AP3chim_-T157L-Q164L (**E**) to a DNA probe containing two CArG-boxes. Increasing amounts of *in vitro* translated protein were incubated together with constant amounts of DNA probe. *k*_*coop*_ values inferred from this particular measurement are depicted below the helical wheel diagrams.

## DISCUSSION

### Conserved leucine residues in the K-domain of SEP3 are required for tetramer formation

Tetramer formation among MIKC-type MADS-domain transcription factors is of central importance for flower development (Theißen and Saedler 2001; Immink, et al. 2009; Melzer and Theißen 2009; Smaczniak, et al. 2012; Theißen, et al. 2016). However, knowledge about the molecular determinants facilitating tetramer formation remains scarce. Our data indicate that substitution of leucines in the K-domain of SEP3 did almost invariably lead to a strong reduction in tetramer formation abilities (Figure 2). This was expected for leucine to proline substitutions within the helical regions of the K-domain as proline has helix-breaking properties. However, also the rather conservative substitution from leucine to alanine in the tetramerization interface (L164A) affected cooperative binding and tetramerization strongly. Similar results have been obtained for substituting other leucine residues in the tetramerization interface by alanine (Puranik, et al. 2014).

The question arises as to why specifically leucine residues are favoured over other hydrophobic amino acid residues. The tetramerization interface forms coiled-coils and it is well established that complex ‘knobs-into-holes’ side chain interactions within the hydrophobic core determine the strength of the interaction between coiled-coils (Parry, et al. 2008). Numerous studies on energetic contributions of different hydrophobic amino acids inside the hydrophobic core revealed that β-branched amino acids (e.g. isoleucine or valine) as well as amino acids with small side chains (e.g. alanine) in heptad repeat ‘d’ positions have a strong destabilizing effect on formation of parallel dimeric coiled-coils (Zhu, et al. 1993; Takei, et al. 2006). The local stereochemical environment at heptad repeat ‘d’ positions instead strongly favours γ-branched amino acids for intermolecular interactions, making leucines uniquely suited at these sites (Zhu, et al. 1993; Betz, et al. 1995; Moitra, et al. 1997; Takei, et al. 2006). This is in line with the observation that L145, which is located at a heptad repeat ‘d’ position but according to structural data not involved in intermolecular interactions (Puranik, et al. 2014) can be mutated to alanine without a decrease in tetramer formation capabilities. In contrast, L164 (also at a heptad repeat ‘d’ position but involved in intermolecular interactions) mutation to alanine leads to a strong decrease in tetramerization. In addition, L145 is by far not as conserved as leucines involved in interactions (Supplementary Figure S8).

A decrease in tetramer formation was also observed for substitution of leucines in the kink region between the two helices, where an effect on helix formation was not predicted (Supplementary Figure S1). However, although the leucine residues in the kink are not directly involved in tetramer formation, they interact intramolecularly with each other to stabilize the kink and thus bring the tetramer interface in a favourable position for protein-protein interactions (Puranik, et al. 2014). It is likely that substitutions to proline or alanine in the kink region altered or destabilized the orientation of the tetramerisation interface and thus impeded tetramer formation indirectly. Similar to the leucines at interacting sites within the helical regions of the K-domain stereochemical restrictions may also in this case favour leucines over other hydrophobic amino acids. This may explain why the L115A mutation in the kink region, which presumably only affects intramolecular interactions, caused a decrease in tetramer formation capabilities.

Taken together, these findings indicate that inter- and intramolecular hydrophobic interactions specifically among leucines are of critical importance for SEP3 homotetramerization. This principle does very likely apply to the entire subfamily of SEP3 proteins, as leucines at interaction positions are evolutionarily highly conserved throughout this subfamily (Figure 4). The evolutionary conserved and important role of leucines is further highlighted by the observation that in the SEP3 ortholog AMtrAGL9 from *A. trichopoda* leucines at positions homologous to those in SEP3 were also of critical importance for tetramer formation (Figure 2). In a recent study Ruelens et al. reconstructed and synthesized the ancestral SEP3 sequence being present at the base of angiosperm evolution (Ruelens, et al. 2017). Interestingly the reconstructed sequence carries all leucine residues that constitute interaction sites within the K-domain of SEP3. In accordance to the presence of the leucine residues the ancestral SEP3 sequence seems to be capable of forming floral quartet-like complexes and mediates the interaction of other ancestral floral homeotic proteins (Ruelens, et al. 2017).

### Structural similarity and interaction specificity among the K-domains of MIKC-type proteins

The K-domain is the second highest conserved domain of MIKC-type proteins (the most highly conserved domain is the MADS-domain) (Kaufmann, et al. 2005). Previous structural predictions indicated that the K-domain is forming coiled-coils in most if not all MIKC-type proteins (Ma, et al. 1991; Riechmann and Meyerowitz 1997; Puranik, et al. 2014). Our analyses indeed strongly support this view. The chemical properties of amino acids that are of importance for intra- and intermolecular interactions in SEP3 are conserved in MIKC-type proteins from all of the 14 subfamilies analysed here. This indicates that most K-domains fold in a structure similar to that determined for SEP3 and that residues that are homologous to interacting sites in the SEP3 homotetramer may also constitute intra- and intermolecular contact points in most other protein family members.

However, although the chemical properties of amino acids important for interactions were conserved in subfamilies other than SEP3, their identity was not always. Whereas the vast majority of leucine residues important for intra- and intermolecular interactions is highly conserved within the SEP3 subfamily, leucine residues are observed at a clearly lower frequency in other subfamilies (Figure 4, Supplementary Figure S4). This indicates that, although the overall structure of the K-domain is conserved in all MIKC-type proteins and probably throughout angiosperm evolution, their tetramerization capabilities may vary depending on the presence of leucines on critical interaction sites. For example, AP3 and PI, who do not possess leucines on all inter- and intramolecular contact points, are unable to form tetramers not involving SEP3 (Melzer and Theißen 2009; Smaczniak, et al. 2012). Indeed, the K3-subdomain of AP3, which is not capable of mediating homotetramer formation, gained this ability when placed in the SEP3 protein context and two leucines were introduced (Figure 4). Thus, we hypothesize that leucines at intra- and intermolecular contact points may not only be necessary but also sufficient for tetramer formation of MIKC-type proteins.

### The ability of SEP3-subfamily proteins to act as hub proteins may depends on highly conserved leucines

Intriguingly, the high conservation of leucines in the K-domain of SEP3-subfamily proteins and their importance for homotetramer formation correlates very well with the crucial function of those proteins as hubs within the PPI network controlling flower development. In addition, proteins like AP3 and PI that have less central positions within the interaction network also lack leucines at several positions critical for tetramerization. It thus appears plausible that leucines in SEP3-subfamily proteins are not only important for homotetramer formation but also play a pivotal role in the formation of heterotetrameric complexes. For example, though a lack of leucines in the kink region of many MIKC-type proteins may destabilizes the orientation of the teramerization interface and prevents homotetramer formation, the high structural stability of the K-domain of SEP3-subfamily proteins that is brought about by intramolecular leucine interactions may serve as a scaffold that helps to align the interaction interface of partner proteins and hence facilitate heterotetramer formation.

The pattern of leucines at the tetramerization interface may be explained in a similar manner. Though data on the interaction of leucines at heptad repeat ‘d’ positions with other amino acids at ‘d’ positions in a heteromeric coiled-coil are scarce, data from leucine zippers indicate that beyond leucine-leucine interactions also interactions of leucines with a number of other amino acids are more favourable than most other interactions not involving any leucine (Fong, et al. 2004).

Taken together, we propose that the leucine residues in SEP3-subfamily proteins serve to facilitate heterotetrameric interactions while at the same time the absence of leucines in the interaction partners prevents homotetramer formation or formation of heterotetramers not involving SEP3-subfamily proteins. This way, tetramerization of many MADS-domain TFs depends on the presence of SEP3-subfamily proteins and probably cannot occur without them.

### Conclusion and outlook

We previously proposed that the dependence of other MIKC-type proteins on SEP3 and LOFSEP-subfamily proteins for tetramer formation facilitated the concerted development of the different floral organs and the evolution of the flower as a single reproductive entity (Melzer, et al. 2014). The evolutionary conservation of leucines in the SEP3 subfamily as opposed to most other subfamilies may be thus one important molecular mechanism that fostered the evolution of the flower.

Importantly, however, coiled-coil interactions are very complex, with amino acids occupying heptad repeat ‘a’, ‘d’, ‘e’ and ‘g’ position playing key roles in determining the affinity and specificity of an interaction (Mason and Arndt 2004; Mason, et al. 2009; Potapov, et al. 2015) and we are far from completely understanding the implications of sequence variations on the different positions for MIKC-type protein interactions. For example, polar and charged residues are observed at heptad repeat ‘d’ positions in a number of MIKC-type protein subfamilies and those would be expected to not only hinder homotetramerization but also heterotetramerization with SEP3-subfamily proteins. Furthermore, subfamily specific patterns of charged residues at heptad repeat ‘e’ and ‘g’ positions can be observed that would be expected to contribute to interaction specificity. These charge distribution patterns could probably explain why heterotetramers are usually formed in favour of homotetramers. Although our findings bring us one step closer towards solving the code for floral quartet-like complex formation, additional structural and biophysical analyses are required to more completely understand the molecular mechanisms and evolutionary patterns of MIKC-type protein interactions. This will eventually also lead to a better understanding as to why this transcription factor family expanded in seed plants and plays a role in virtually every reproductive developmental process.

## METHODS

### Cloning procedures and site-directed mutagenesis

The plasmids for *in vitro* transcription/translation of *SEP3* (NCBI accession: NM_102272), *AP3* (NM_115294), *PI* (NM_122031) and *AMtrAGL9* (KF925502) (pTNT-SEP3, pSPUTK-AP3, pSPUTK-PI and pSPUTK-AMtrAGL9) have been generated previously (Melzer, et al. 2009; Melzer, et al. 2014). The cDNA sequences for the single- and double amino acid substitution mutants of SEP3 were created by site-directed mutagenesis PCR according to the Q5 Site-Directed Mutagenesis Manual (New England Biolabs). The cDNA sequence for the chimeric protein SEP3_AP3chim_ was created by megaprimer-mediated mutagenesis PCR for domain substitutions according to (Perez, et al. 2006).

### Design of DNA probes and radioactive labeling

Design and preparation of DNA probes have been described previously (Melzer, et al. 2009). The CArG-box sequence 5’-CCAAATAAGG-3’ that was used for all DNA probes was derived from the regulatory intron of *AGAMOUS*. For studies on homotetramer formation a 151 nt long DNA probe was used that contained two CArG-boxes in a distance of 63 bp, i.e. 6 helical turns (sequence: 5’-TCGAG GTCGG AAATT TAATT ATATT <u>CCAAA TAAGG</u> AAAGT ATGGA ACGTT CGACG GTATC GATAA GCTTG ATGAA ATTTA ATTAT ATT<u>CC AAATA AGG</u>AA AGTAT GGAAC GTTAT CGAAT TCCTG CAGCC CGGGG GATCC ACTAG TTCTA G -3’, CArG-box sequences are underlined). Saturation binding assays to quantify dimer binding affinities were performed with a 51 nt long DNA probe harboring a single CArG-box in the center (sequence: 3’-AATTC GAAAT TTAAT TATAT T<u>CCAA ATAAG G</u>AAAG TATGG AACGT TGAAT T-5’, CArG-box sequence is underlined). The DNA probes were radioactively labeled via Klenow fill-in reaction of 5’-overhangs with [α-^32^P] dATP.

### *In vitro* transcription/translation and electrophoretic mobility shift assay

Proteins were produced *in vitro* using the TNT SP6 Quick Coupled Transcription/Translation System (Promega) according to the manufacturer’s instructions and used directly without freezing and thawing. The composition of the protein-DNA binding reaction buffer was essentially as described (Egea-Cortines, et al. 1999), with final concentration of 1.6 mM EDTA, 10.3 mM HEPES, 1 mM DTT, 1.3 mM spermidine, 33.3 ng/µl Poly dI/dC, 2.5 % CHAPS, 4.3 % glycerol, and a minimum of 1.3 µg/µl BSA. The amounts of protein, DNA probe and BSA were varied according to the performed assay. For cooperative DNA-binding studies to infer tetramer formation capabilities a constant amount of 0.1 ng of a DNA probe containing two CArG-boxes in a distance of six helical turns was co-incubated with variable amounts of *in vitro* translated protein ranging from 0.05 µl to 3 µl. Variable amounts of applied *in vitro* translated protein were compensated by adding appropriate volumes of BSA (10 µg/µl). For saturation binding assays to quantify dimer binding affinities a constant amount of 2 to 5 µl *in vitro* translated protein was co-incubated with variable amounts of a DNA probe containing one CArG-box in the center, ranging from 0.05 to 32 ng as previously described in (Jetha, et al. 2014). Binding reactions had a total volume of 12 µl, were incubated overnight at 4°C and subsequently loaded on a polyacrylamide (5 % acrylamide, 0.1725 % bisacrylamid) 0.5x TBE gel that has been pre-run for 30 min. The gel was run with 0.5x TBE buffer for 2.5 h at 7.5 V/cm and afterwards dried and exposed onto a phosphorimaging screen to quantify signal intensities.

### Quantification of cooperative DNA-binding

For each lane of the EMSA gel relative signal intensities of all fractions were measured using Multi Gauge 3.1 (Fujifilm). The equations that were used to quantify the ability for cooperative DNA-binding of two dimers to a DNA probe carrying two CArG-boxes have been described previously (Senear and Brenowitz 1991; Melzer, et al. 2009). Briefly, if the relative concentration of unbound DNA probe [Y_0_] (signal of high electrophoretic mobility), a DNA probe bound by two proteins [Y_2_] (signal of intermediate electrophoretic mobility), and a DNA probe bound by four proteins [Y_4_] (signal of low electrophoretic mobility) are described as a function of applied protein [P2],

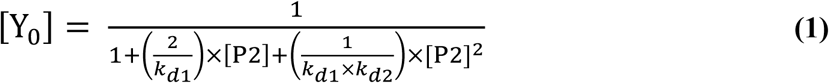

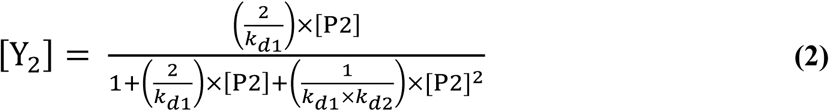

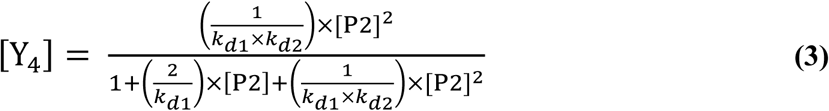

then *k*_*d1*_ is the dissociation constant for binding of a protein dimer to a DNA probe with two unoccupied binding sites and *k*_*d2*_ is the dissociation constant for binding of a second protein dimer to a DNA probe where one of the two binding sites is already occupied. By nonlinear regression of the measured signal intensities of the three fractions to equation (1) to (3), *k*_*d1*_ and *k*_*d2*_ were estimated using GraphPad Prism 5 (GraphPad Software). As we used *in vitro* transcription/translation for protein production, the exact protein concentrations were unknown. Therefore the amount of applied *in vitro* transcription/translation mixture was used as proxy for [P2], as previously described (Melzer, et al. 2009). As a result of the unknown protein concentrations the estimated values for *k*_*d1*_ and *k*_*d2*_ depend on the *in vitro* transcription/translation efficiency and can only be considered as relative values. However, estimating a cooperativity constant *k*_*coop*_ (defined as the ratio of *k*_*d1*_ and *k*_*d2*_) is still possible:

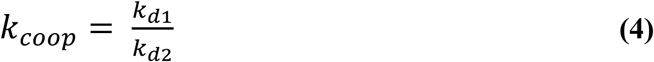

As described earlier, *k*_*coop*_ values of ≈ 200 were the upper limit that could be determined with our experimental setup (Jetha, et al. 2014).

#### Saturation binding assay

To estimate the dissociation constant for binding of a protein dimer to a single DNA-binding site *k*_*d*_, saturation binding assays with a DNA probe carrying a single CArG-box were performed. The equation that was used to infer *k*_*d*_ has been described previously (Jetha, et al. 2014). *k*_*d*_ can be defined as

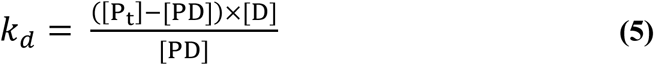

with [PD], [P_t_], and [D] being the concentration of the protein-DNA complex, total protein, and unbound DNA probe, respectively. By expressing [PD] as a function of [D] for increasing concentrations of applied DNA probe, [P_t_] and *k*_*d*_ were determined via nonlinear regression using GraphPad Prism 5.

### Multiple sequence alignments and *in silico* sequence analysis

For analyses on amino acid preferences of different MIKC-type MADS-domain protein subfamilies throughout the K-domain a comprehensive sequence collection was compiled. Via BLAST search (Altschul, et al. 1990) representatives of all 14 subfamilies (Becker and Theißen 2003; Gramzow and Theißen 2013) of MIKC-type proteins present in *A. thaliana* (AP1-, AP3-, PI-, AG-, ABS-, SEP3-, LOFSEP-, AGL6-, AGL12-, AGL15-, AGL17-, FLC-, TM3-, and SVP-subfamily) were collected using the amino acid sequences of *A. thaliana* AP1, AP3, PI, AG, ABS, SEP3, SEP1, AGL6, AGL12, AGL15, AGL17, FLC, SOC1, and SVP, respectively, as query. To cover a broad set of species, six individual searches were performed for each subfamily. Each of those searches was restricted to a different group of seed plants: core eudicots, early diverging eudicots, monocots, magnoliids, early diverging angiosperms, and gymnosperms. For sequences from core eudicots the search queries were restricted to asterids (BLAST tax-ID: 71274), Dilleniaceae (24942), Caryophyllidea (108240), Santalales (41947), Berberidopsidales (403664), Saxifragales (41946), rosids (71275), and Gunnerales (232382); for sequences from early diverging eudicots the search queries were restricted to Proteales (232378), Buxales (280577), and Ranunculales (41768); for sequences from monocots and magnoliids, respectively, the queries were restricted to the corresponding predefined organism groups implemented in BLAST (tax-ID: 4447 and 232347, respectively); for sequences from early diverging angiosperms the queries were restricted to Austrobaileyales (82956), Hydatellaceae (178426), Nymphaeales (261007), and Amborella (13332); and for sequences from gymnosperms the queries were restricted to Gnetales (3378), Pinaceae (3318), Taxaceae (25623), Cephalotaxus (50178), Cupressaceae (3367), Araucariaceae (25664), Podocarpaceae (3362), Ginkgoales (3308), and Cycadales (3297). For each of the 84 resulting BLAST searches the amino acid sequences of all hits were downloaded (if more than 100 sequences were found, only top 100 hits according to the total score calculated by BLAST were downloaded). The results of all BLAST searches were combined into a single data set, all completely redundant sequences as well as all sequences that did not constitute MIKC-type proteins were removed and the remaining sequences were aligned with Mafft applying E-INS-i mode using Jalview (Waterhouse, et al. 2009; Katoh and Standley 2013). The subfamily assignment of each sequence was performed according to its clustering within a phylogenetic tree calculated with MrBayes (based on MADS-, I- and K-domain sequences, applying mixed AA model with 20 million generations, 50% burn-in, and a sample frequency of 1000) (Huelsenbeck and Ronquist 2001). All sequences with uncertain subfamily assignment were removed. To optimize the alignment quality of the K-domain 133 further sequences were removed that produced gaps and that did not appear to be representative for the respective subfamily. The final sequence collection comprised 1325 MIKC-type protein sequences.

Relative sequence similarities at homologous sites were calculated with R (https://www.R-project.org/). Each pair of amino acids at equivalent sites were assigned a similarity score based on BLOSUM40 values that were normalized to 1 via dividing by the maximum value of the respective amino acid. Subsequently, all pairwise similarity scores were averaged to calculate the mean relative sequence similarity for all amino acid positions within the K-domain. BLOSUM40 was chosen because the average sequence identity within the K-domain of all examined sequences was about 40 %. Box-plots and line graphs of sequence similarity scores were created with SPSS (IBM). Statistical significance of sequence similarity differences were tested via Mann-Whitney U-tests implemented in SPSS.

Subfamily specific amino acid frequencies and mean hydrophobicity values for positions within the K-domain were calculated with R. SEP3 K-domain crystal structure pictures were created with Swiss-PdbViewer (SIB). Helical wheel diagrams were created with R. Coiled-coil predictions to preselect potential positions for single and double amino acid substitutions were performed with COILS (Lupas, et al. 1991).

## ACKNOWLEDGEMENTS

We are grateful to Fredo-Torpedo (Fred Ferber), Chris-Master (Christian Gafert) and Tanja Schulze for valuable help with some experiments. This work was supported by the DFG (Deutsche Forschungsgemeinschaft) grant to G.T. and R.M. (TH417/5-3). R.M. received a post-doctoral fellowship from the Carl Zeiss Foundation.

